# Rapid Retreat of the Pacific Maritime Forest

**DOI:** 10.1101/2020.08.31.273847

**Authors:** George Kral, Melodie Putnam, David Rupp

## Abstract

The temperate maritime climate of the Pacific Northwest region of the United States supports the world’s tallest and most economically productive conifer forests. These forests are vital to local ecosystems and society, and climate perturbations are likely to adversely affect the services these forests provide. This study presents a simple, easily replicated methodology for assessing effects of climate change in these local forests, using species with differential climatic ranges as ecological barometers. A comparative analysis of warm-adapted and cool-adapted species co-occurring within a warming but otherwise climatically homogenous area near the southeast margin of the Pacific maritime forest reveals dramatic differences in tree health and mortality between these climatically differentiated species groups. Our results strongly suggest a rapid decline at the southeastern extent of the Pacific maritime temperate forest, and a need to immediately modify local land management practices to address this new reality.

## Introduction

Temperate and boreal forests are economically and ecologically vital. The temperate maritime forests of northwestern North America are of particular economic importance; the Pacific Slope forests of Oregon, Washington and northern California alone contain a third of the total softwood volume of the United States (Oswalt 2014). In Canada, approximately 40 percent of all timber produced in 2018 came from Pacific Slope forests (Natural Resources Canada 2018). Because of their ecological and economic importance, there has been much conjecture regarding potential response of these and other temperate forest systems to climate change (Daniels 2011, Hamann 2006, Vose 2012, Terrier 2013, HilleRisLambers 2015, Carnicer 2013, Keane 2001).

While numerous studies predict or document forest declines and die-off events in the arid west (Allen and Breshears 1998, Breshears 2005, Adams 2009, Allen 2010, Smith 2015, Clifford 2013, Goulden 2019), there have been relatively few studies investigating actual climate-associated forest responses in temperate systems, and the findings of these studies have been mixed. Some studies indicate a climate change influence on temperate forests. Monleon (2015), for instance, found modest but significant overall tendencies toward upward and northerly range shifts of tree taxa on the western coast of North America by comparing differential elevational and latitudinal distribution of mature trees and their progeny, although some of the evidence was equivocal or even contradictory. Looking at the entire eastern United States, Woodall (2009) identified significant northerly migration trends in northerly-ranging versus southerly-ranging species, with northward migration rates estimated as high as 1 km per year. In a study of 76 long-term forest plots in the western US, van Mantgem (2009) found significant increases in tree mortality across all species in California, the Pacific Northwest, and the interior West.

A subsequent investigation in western Washington, however, found no significant increase in tree mortality looking at some of the same forest types (Acker 2015). Monleon (2015), Woodall (2009) and other similar studies rely on snapshot differences in the distribution of seedlings and saplings versus mature trees as evidence of shifting ranges. The assumption that these differences are climate-driven is challenged by Máliš (2016) who presents evidence that the offset ranges of tree progeny are more driven by ontogenetic effects, such as differential niche occupancy between seedlings and mature trees, than by changing climate. Leak (2012) found no evidence of range shifts in long-term forest plot data of easten hemlock (*Tsuga canadensis*) and red spruce (*Picea rubens*), or in upslope seedling recruitment of several species of hardwoods and conifers in the northeastern US. Generally, observational data have suggested that shifts in extent and composition of temperate forests have been slow to occur in the absence of acute disturbance, and some researchers have concluded that increased temperature is unlikely to be a singular driver in tree species decline and migration in temperate forests, at least in the near term (HilleRisLambers 2015, Leak 2012).

Even in temperate forests, however, increasing temperatures may exacerbate the effects of periodic drought, insects and pathogens. Synergistic interactions of these factors have been implicated in a number of cases of widespread tree die-offs in more arid regions (Breshears 2005, Adams 2009, Smith 2015, Goulden 2019). These and other studies of tree die-offs and forest declines are congruent with the hypothesis that increasing temperatures and severity of periodic drought could erode the southerly margins of northern hemisphere temperate forests.

The presence of a particular species provides direct evidence that climatic conditions at the time of its establishment at a given locality were within the tolerances of the species as a whole. Likewise, a species decline at that locality suggests that one or more environmental variables have moved beyond that species’ tolerance. Individually, one species’ decline or local extinction provides limited circumstantial evidence of any particular environmental change. Multiple species in decline, however, present an opportunity to analyze commonalities in declining species’ habitat requirements. These kinds of observations and analyses can identify local impacts of environmental change and provide evidence-based guidance for model refinement and on-the-ground decision making.

As an example, striking levels of tree decline and mortality in the Willamette Valley, Oregon, U.S., are readily visible and of growing concern to foresters (Withrow-Robinson 2018, Oregon Department of Forestry 2019). Tree decline and mortality have become apparent only in certain species, however, and not others (Fig. 1). In this study, we examine the condition of four temperate maritime species and four Mediterranean climate-adapted species co-occurring in the Willamette Valley. We sampled 674 individual trees in 55 stands and assessed the health of each tree. We also cultured tissue samples from declining black hawthorn trees to rule out acute pathogenic causation for observed decline in this species, a decline which has not previously been investigated. We analyzed and compared the current condition of northerly-ranging and southerly-ranging tree species to determine if patterns emerged relative to species ranges.

**Figure 1:**
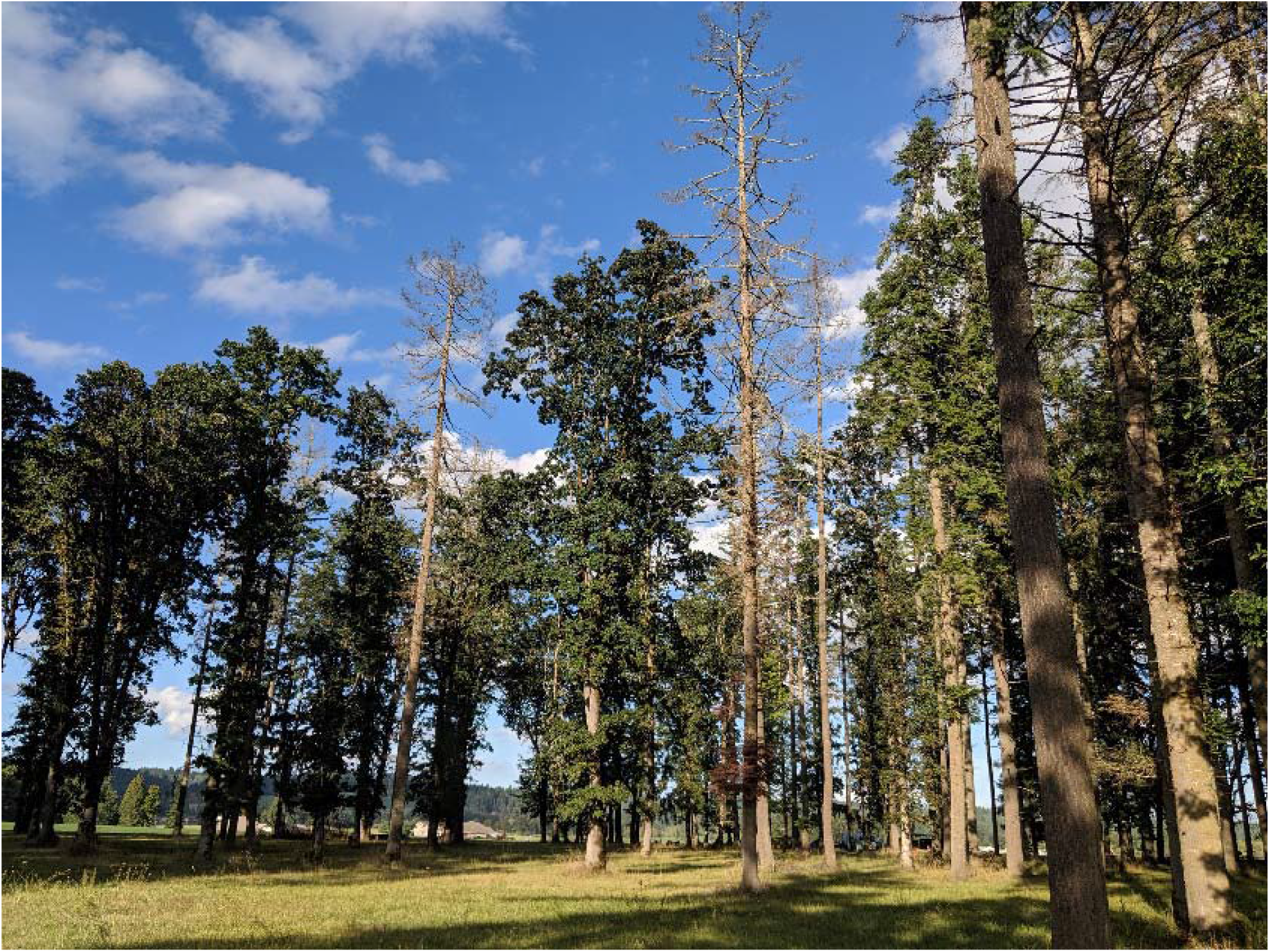
Mortality of grand fir in the Willamette Valley, Oregon in 2019. Note adjacent healthy Garry oaks.

## Methods

### Study System

As a model system for investigating climate impacts in forests, the Willamette Valley presents an opportunity for side-by-side performance evaluations of northward-ranging and southward-ranging species in a warming climate. The Willamette Valley is transitional between the temperate maritime climate zone to the north and the Mediterranean climate to the south, and elements of both temperate and Mediterranean plant communities can be found here, growing together (OregonFlora 2020).

The climate of the Willamette Valley has become markedly warmer in recent decades compared to most of the previous century, consistent with the larger regional pattern (Abatzoglou et al., 2014; Oregon Climate Change Research Institute 2019). Since 1990, especially acute temperature increases have emerged in mid and late-summer, coincident with an intensification of the annual summer dry season.

Examining the records from the three stations that latitudinally span the Willamette Valley floor reveals a clear pattern of recent elevated summer temperatures compared to a 1920-1990 baseline (Figure 2) when most natural forest stands in the Willamette Valley became established, including all but one of the stands surveyed in this investigation. Averaging over the three stations, the five-year mean summer (July-August) temperature for both 2002-2006 and 2014-2018 was about 1.7°C above baseline. As a measure of how extreme this anomaly is, 1.7°C amounts to 2.7 standard deviations of the 1920-2019 five-year moving average of summer anomalies. At the annual-scale, the 3-station mean summer temperature was below seasonal norm only three times in the last three decades (1990-2019), and of the eleven warmest summers since 1920, four were since 2014 (2014, 2015, 2017, and 2018).

**Figure 2.**
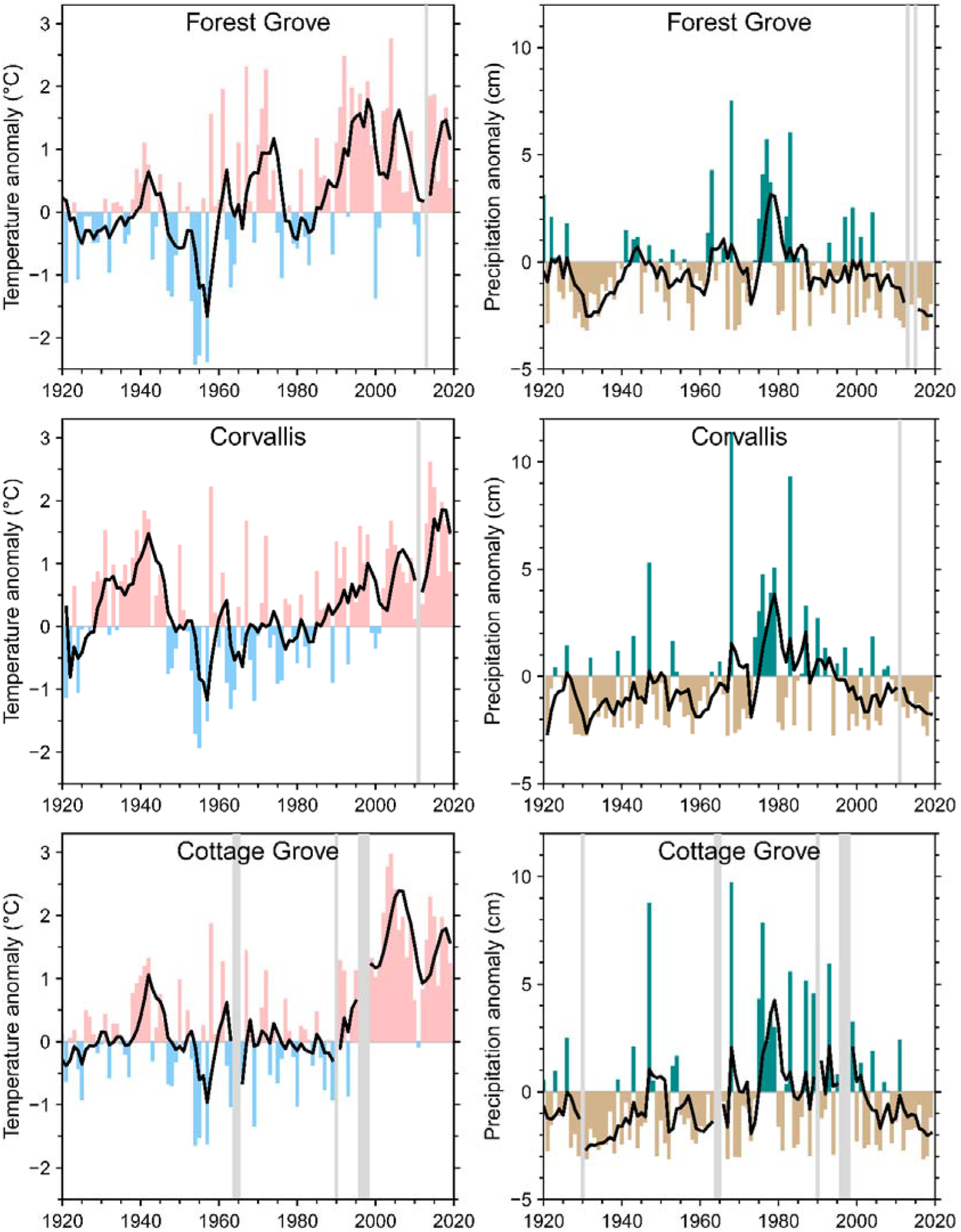
Left column: July-August mean temperature anomalies from a 1940-1990 baseline period at Forest Grove (station USC00352997), Corvallis (station USC00351862), and Cottage Grove (station USC00351897), Oregon. The black line shows the 5-year backward moving average of the temperature anomalies. Right column: Same as left column but for July-August total precipitation anomalies. July-August periods with more than 4 days of missing data were excluded (gray shading). Data source: Global Historical Climate Network (Menne et al., 2012)

Along with elevated temperatures, persistent deficits in summer precipitation have emerged since 2000 (Figure 2). July-August rainfall averaged over the same three stations during this period was 60% of normal (1940-1990 baseline). It is worth noting that from 1940 to 1946 and 1951 to 1964, the 5-year moving average of July-August precipitation was also below average. Unlike the recent long-term summer meteorological drought, however, over most of this earlier period, below-average summer precipitation was associated with near or below-average summer temperature.

The combination of high temperature and low humidity can be expressed through high vapor pressure deficit (VPD). High VPD can stress trees and, when high enough, lead to hydraulic failure and mortality (Bréda et al., 2006; Eamus et al., 2013; McDowell et al., 2008). In the Pacific Northwest, high VPD has been associated with recent decreased growth of Douglas fir (*Psuedotsuga menziesii*) (Restaino et al., 2016). Reliable estimates of VPD for the Willamette Valley at a small number of stations can be made back only to 1948 based on (Daly et al., 2015) and those data include some substantial gaps. Despite these limitations, estimates of July-August mean daily maximum VPD (VPDmax) at three stations that latitudinally span much of the Willamette Valley indicate a prolonged period of above-average summer VPDmax since 2002 relative to a 1948-1990 baseline (Figure 3). VPDmax was particularly high during several years of the last decade; the years with the highest summer VPDmax since 1948 were 2015, 2018, 2017 and 2014 (in descending order), averaged over the three stations. The five-year moving averages of these data indicate the kind of persistent heat and drought stress which Buotte (2018), Daniels (2011) and others have linked with decreased allocation of tree resources to growth, tree decline and increased tree mortality.

**Figure 3.**
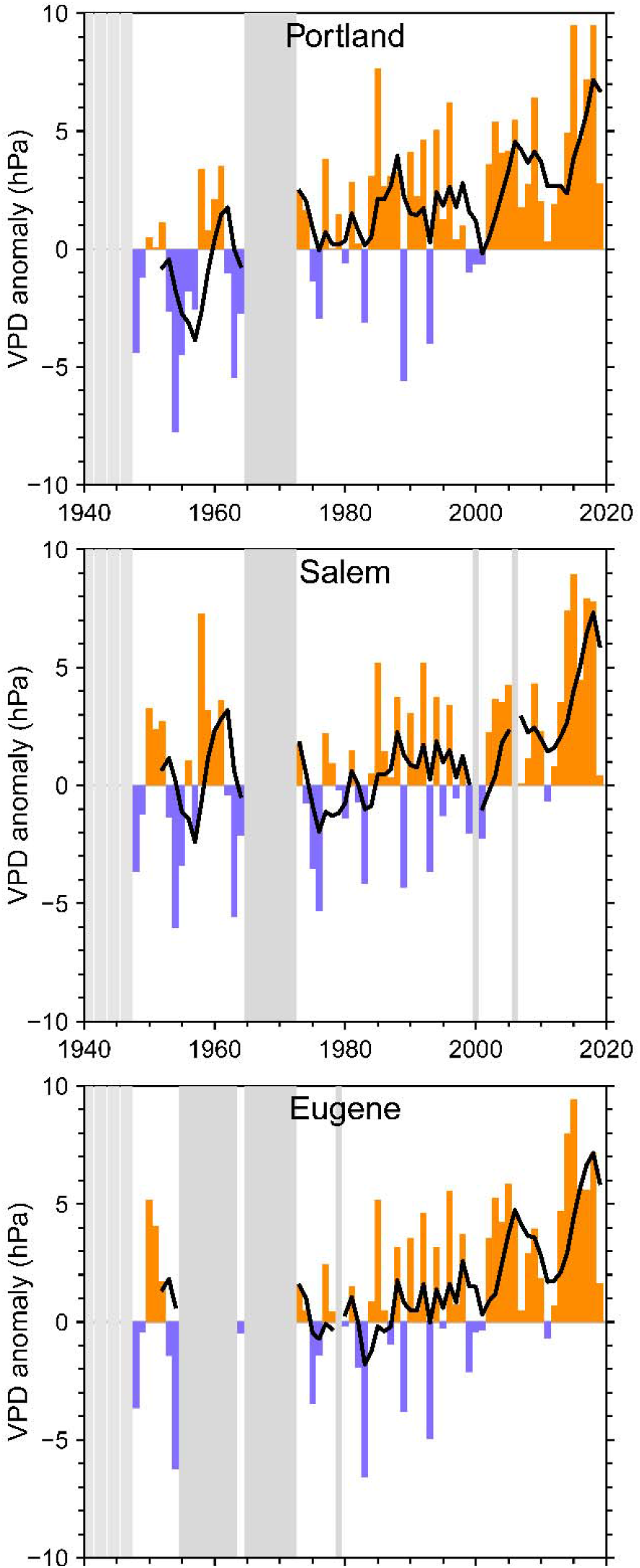
July-August mean of daily maximum vapor pressure deficit anomalies from 1940-1990 baseline period at Portland (station KPDX), Salem (station KSLE), and Eugene (station KEUG), Oregon, airports The black line shows the 5-year backward moving average of the anomalies. July-August periods with more than 4 days of invalid data were excluded (gray shading).

Concurrent with the prolonged and ongoing period of warming and, over the last two decades, the intensification of summer drought, there has been large-scale tree mortality throughout the Willamette Valley (Withrow-Robinson 2018, Buhl 2017). Symptoms of tree decline have appeared in certain tree taxa, however, and not others. Some of the first trees to exhibit widespread decline were black hawthorn (*Crataegus gaylussacia*). Necrotic tissue from dead and dying hawthorns submitted to the Oregon Department of Forestry for pathogen analysis between 2006-2008 exhibited no identifiable acute pathogens at that time (Alan Kanaskie, Oregon Department of Forestry, unpublished data). Since 2010, observers have identified several other woody taxa showing signs of decline in the Willamette Valley, including western redcedar (*Thuja plicata*), grand fir (*Abies grandis*), red alder (*Alnus rubra*) and others (Oregon State University Extension Service 2018, Oregon Department of Forestry 2019, Sims 2015). All of these declining species share similar distributions in areas with cool, moist, moderate climates. They range from the Alaska panhandle or western British Columbia southward through western Montana, northern Idaho, western Washington and Oregon, and coastal California (Fig. 4, right panel). Indicative of the range of these species are narrow extensions southward along the northern and central California coastal strip, where historically, summer temperatures have been reliably cool. They are rare or absent in the Columbia Basin, interior Klamath region, the Sierra Nevada foothills and other interior portions of California (Consortium of Pacific Northwest Herbaria 2020, CalFlora 2020).

**Fig 4:**
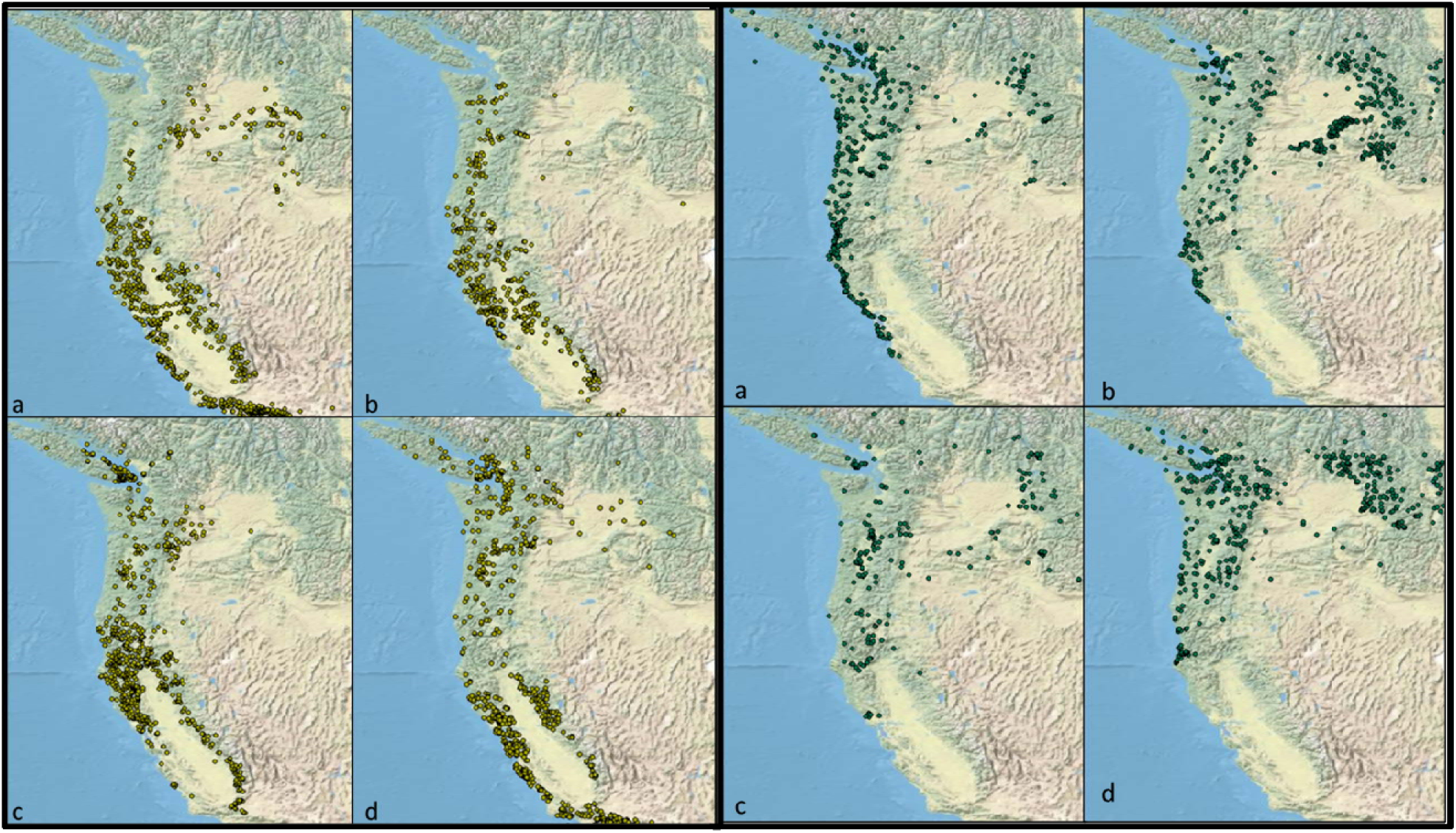
Distribution of four Mediterranean climate-adapted species:LEFT (a) <u>Alnus rhombifolia</u> (white alder) (b) <u>Fraxinus latifolia</u> (Oregon ash), (c) <u>Quercus garryana</u> (Garry oak), and (d) <u>Acer macrophyllum</u> (bigleaf maple), and four temperate maritime species RIGHT (a) <u>Alnus rubra</u> (red alder) (b) <u>Abies grandis</u> (grand fir) (c) <u>Crataegus gaylussacia</u>(black hawthorn) and (d) <u>Thuja plicata</u> (western redcedar) (data from CalFlora.org, PNWherbaria.org)

Meanwhile, several tree species with more southerly and interior ranges co-occur in the Willamette Valley with temperate maritime species (Fig. 4, left panel). These more southerly species are performing relatively well and do not exhibit evidence of decline. All of these species share broad ranges in interior portions of California, and also in the comparatively warm and dry Columbia Basin and the Snake River Valley (Consortium of Pacific Northwest Herbaria 2020, CalFlora 2020. Little 1971).

### Stand and Sample Tree Selection\

For all species except white alder, survey stands were selected along north-south and east-west transects through the Willamette Valley (Fig. 5). In some cases, it was possible to survey multiple species within a single survey stand. From the beginning point of each transect, the first stand of each species encountered was surveyed. Following every 16-km increment thereafter along the transect, the first accessible stand was surveyed. For white alder, which has a very sporadic distribution within the study area, sample data was collected from seven study stands broadly distributed throughout the Willamette Valley.

**Figure 5:**
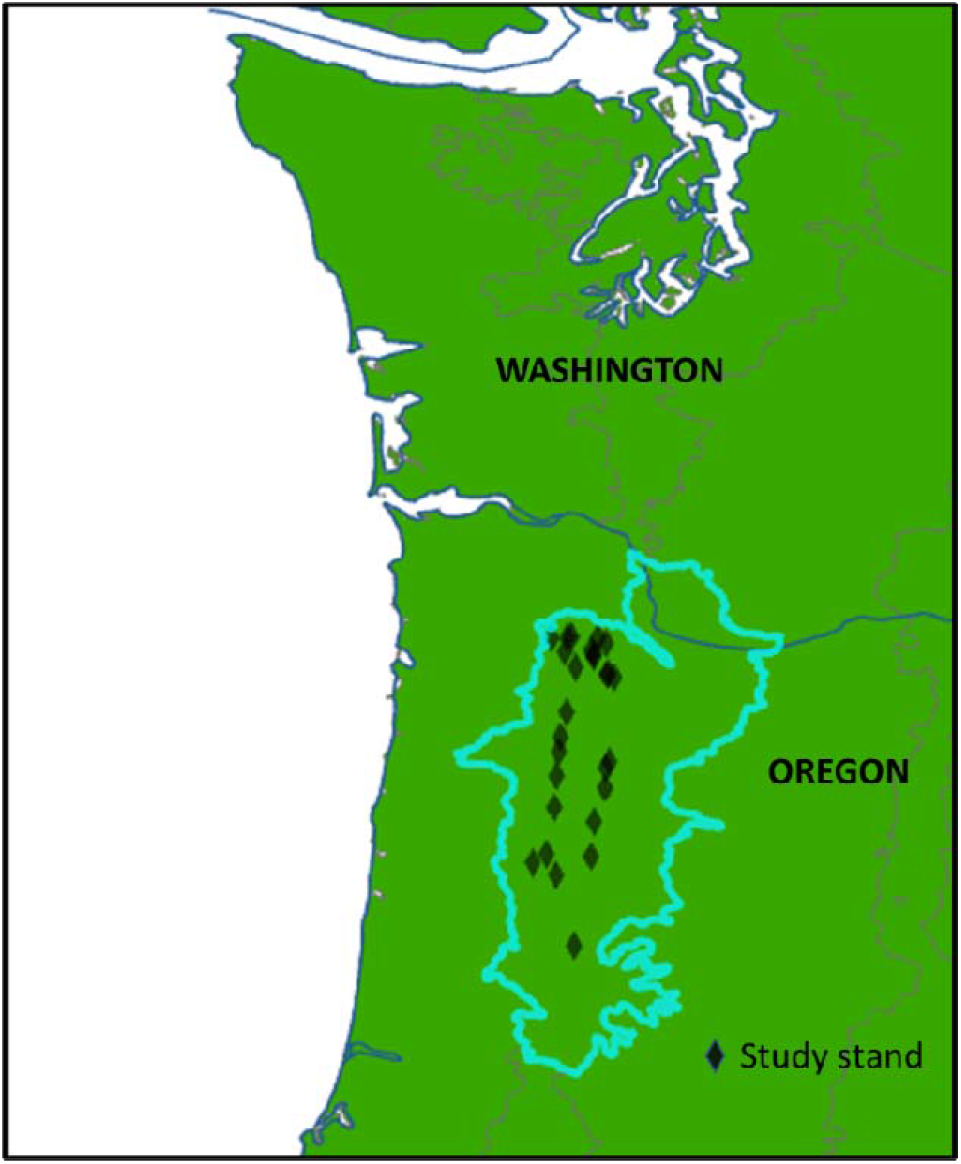
Willamette Valley ecoregion and stand survey locations

Distributional differences among the species required different sample sizes per survey point. Stand-forming species often presented large numbers of individuals from which to select. In these cases, up to 25 trees (10 trees for hawthorn) per sample point were assessed. Only dominant and co-dominant individuals in stands, well-established saplings (minimum diameter 5cm), free-to-grow individuals not suppressed by competing vegetation, and formerly dominant/codominant, recently dead class I snags (characterized by sound sapwood, intact fine branches and tight bark, Cline 1980) were assessed. Trees and snags were selected and assessed simply as they were encountered. Once the maximum sample number of encountered individuals was reached, or all of the available trees had been assessed, the survey for that species was concluded at that survey point.

### Sample Tree Assessment

Each tree was assessed for indicators of crown decline, including top dieback, branch flagging, chlorosis/crown thinning, epicormic branching, and mortality. Other conditions such as bleeding cankers were noted on individual trees. A 0.2-point deduction was assessed for every decline indicator. Perfectly healthy trees with no decline indicators received a score of 1. Dead individuals received a score of 0. Trees were rated as good (score > 0.7), fair (0.7 >= score > 0.4), poor (0.4 >= score > 0), and dead (score = 0).

### Pathogen Investigation

To investigate the possibility of an acute, emerging pathogen in rapidly declining black hawthorn stands, samples of root, trunk or branch tissue from 14 individuals showing moderate to advanced decline systems were evaluated for known pathogenic organisms. Samples were examined visually and microscopically for evidence of fungal or bacterial disease such as branch or trunk cankers, root rot, crown rot, or other symptoms of biotic disease. Select tissue showing distinct and sharp transitions from healthy to necrotic areas and those without such transitional zones were sampled, disinfected in 10% household bleach (1:9 bleach:water) for 3 minutes, rinsed, and air dried under a laminar flow hood until no free moisture was visible. Tissue from canker margins, when present, or from the region nearest the area of dieback when no cankers were present, was sub-sampled and aseptically placed on water agar (1.5 %) and ¼ strength potato dextrose agar containing 100 ppm streptomycin to culture for fungi. Culture plates were incubated at 20 °C for 7-14 days or until sporulation occurred. Fungal cultures were identified based on their morphology.

## Results

### Stand and Tree Condition

Dramatic difference in tree condition and rates of mortality are apparent between temperate maritime and Mediterranean tree species surveyed (Figures 6 and 7). On average, 30% of temperate maritime trees have recently died, and only 30% are in good condition (Figure 8, table 1). By contrast, 94% of the Mediterranean-adapted trees are in good condition and less than 2% have recently died.

**Table 1:** Sampling intensity and tree health analysis of four maritime-adapted and four Mediterranean climate-adapted species

| Species | Trees Sampled | No. Stands | Avg Score | Good | Fair | Poor | Dead |
| --- | --- | --- | --- | --- | --- | --- | --- |
| *Abies grandis* | 75 | 8 | 0.44 | 36% | 12% | 7% | 45% |
| *Alnus rubra* | 100 | 7 | 0.36 | 21% | 16% | 23% | 40% |
| *Crataegus gaylussacia* | 100 | 10 | 0.43 | 24% | 19% | 36% | 21% |
| *Thuja plicata* | 86 | 4 | 0.60 | 37% | 29% | 20% | 14% |
| *Acer macrophyllum* | 75 | 5 | 0.95 | 95% | 4% | 1% | 0% |
| *Alnus rhombifolia* | 85 | 7 | 0.86 | 88% | 6% | 1% | 5% |
| *Fraxinus latifolia* | 75 | 4 | 0.95 | 96% | 1% | 1% | 1% |
| *Quercus garryana* | 78 | 10 | 0.97 | 97% | 0% | 1% | 1% |
| Total | 674 | 55 |  |  |  |  |  |
| temperate species | 361 |  | 0.46 | 30% | 19% | 21% | 30% |
| Mediterranean species | 313 |  | 0.93 | 94% | 3% | 1% | 2% |

**Figure 6:**
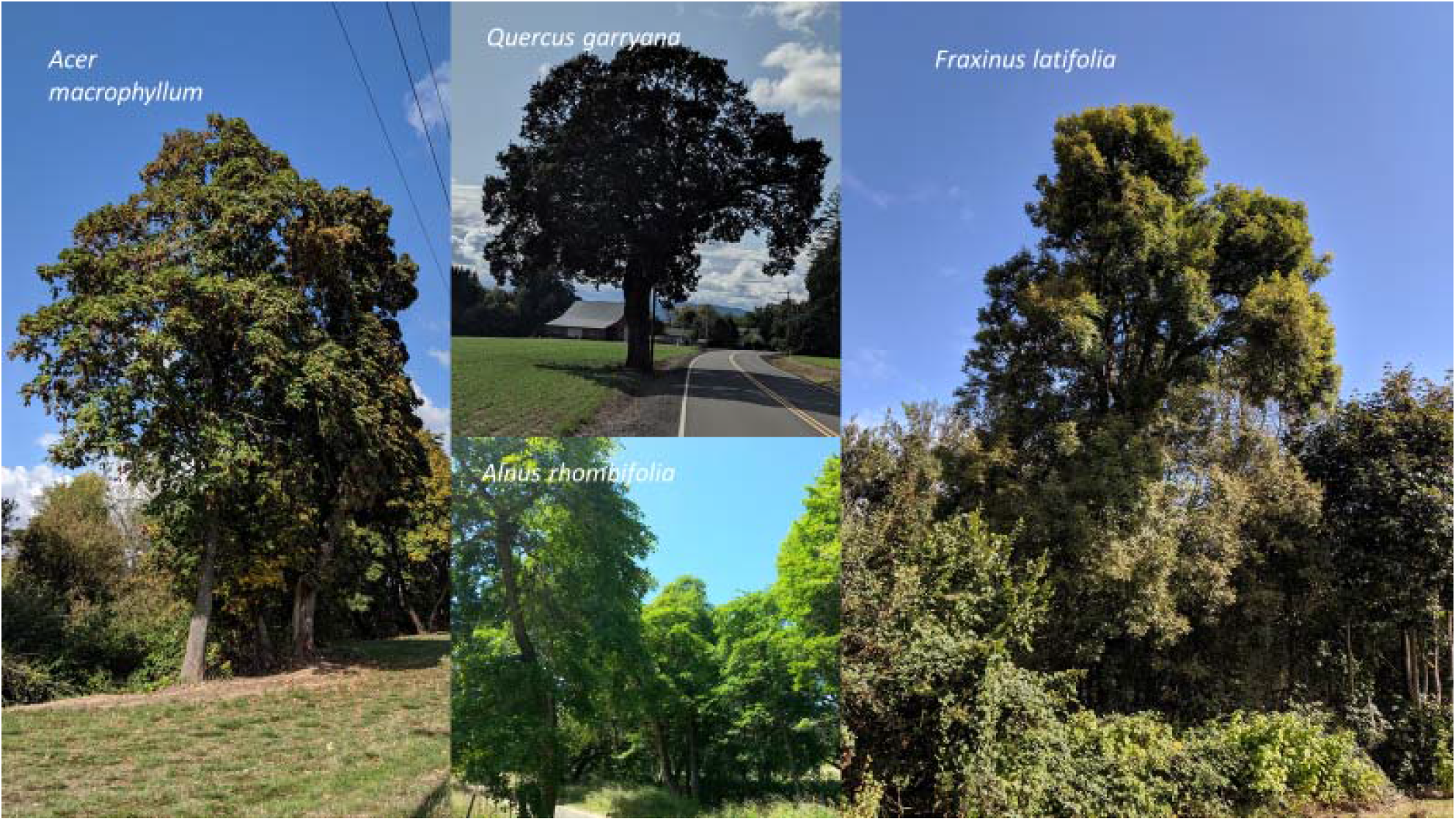
Typical specimens of southerly-ranging species within the study area. Note full, leafy crowns.

**Figure 7:**
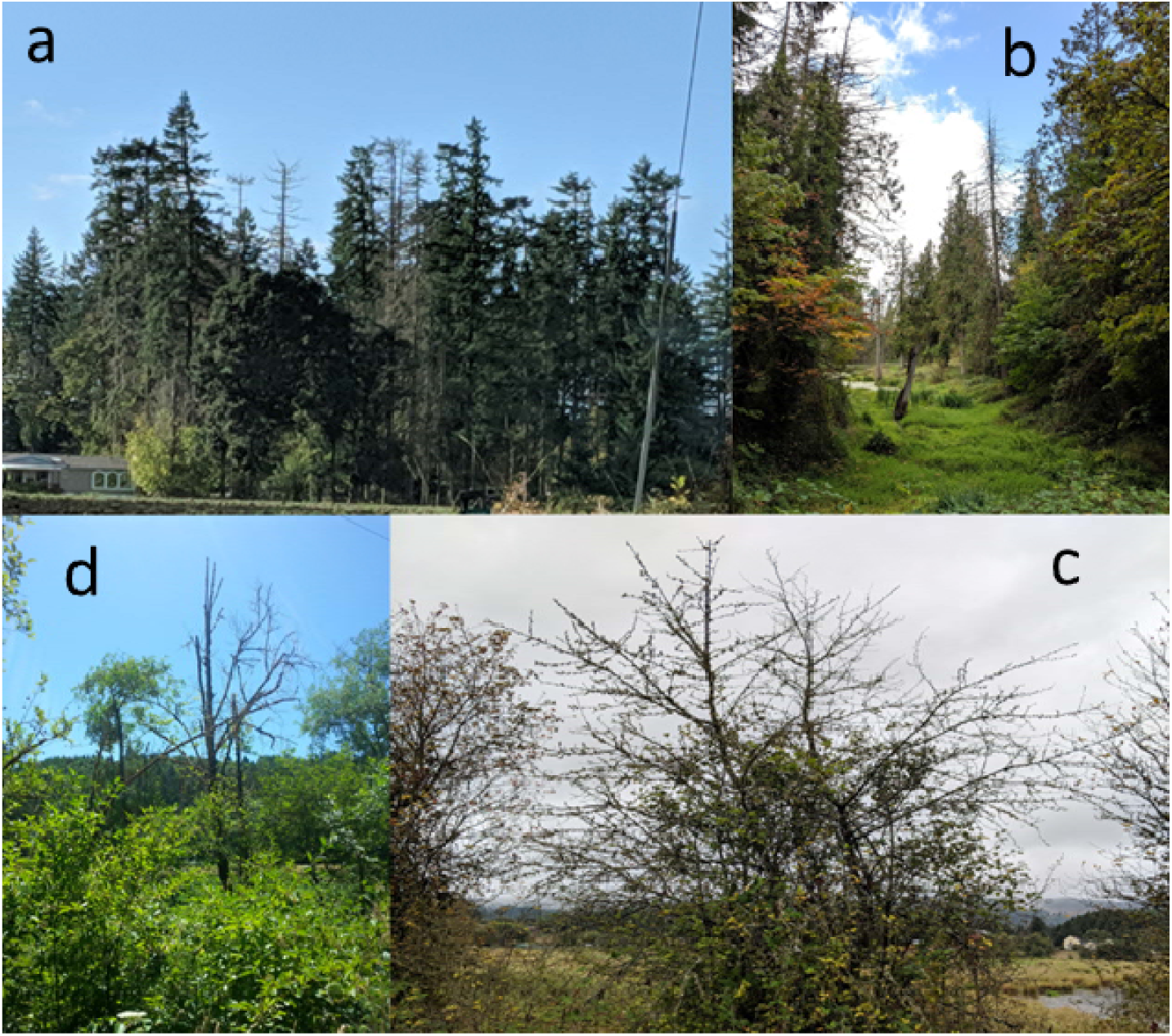
Typical decline in four temperate species. Clockwise from top left (a) grand fir in mixed stand with <u>Pseudotsuga menziesi</u>i and <u>Quercus garryana</u>; (b) Western redcedar with <u>Acer macrophyllum;</u> (c) black hawthorn; and (d) red alder.

**Figure 8:**
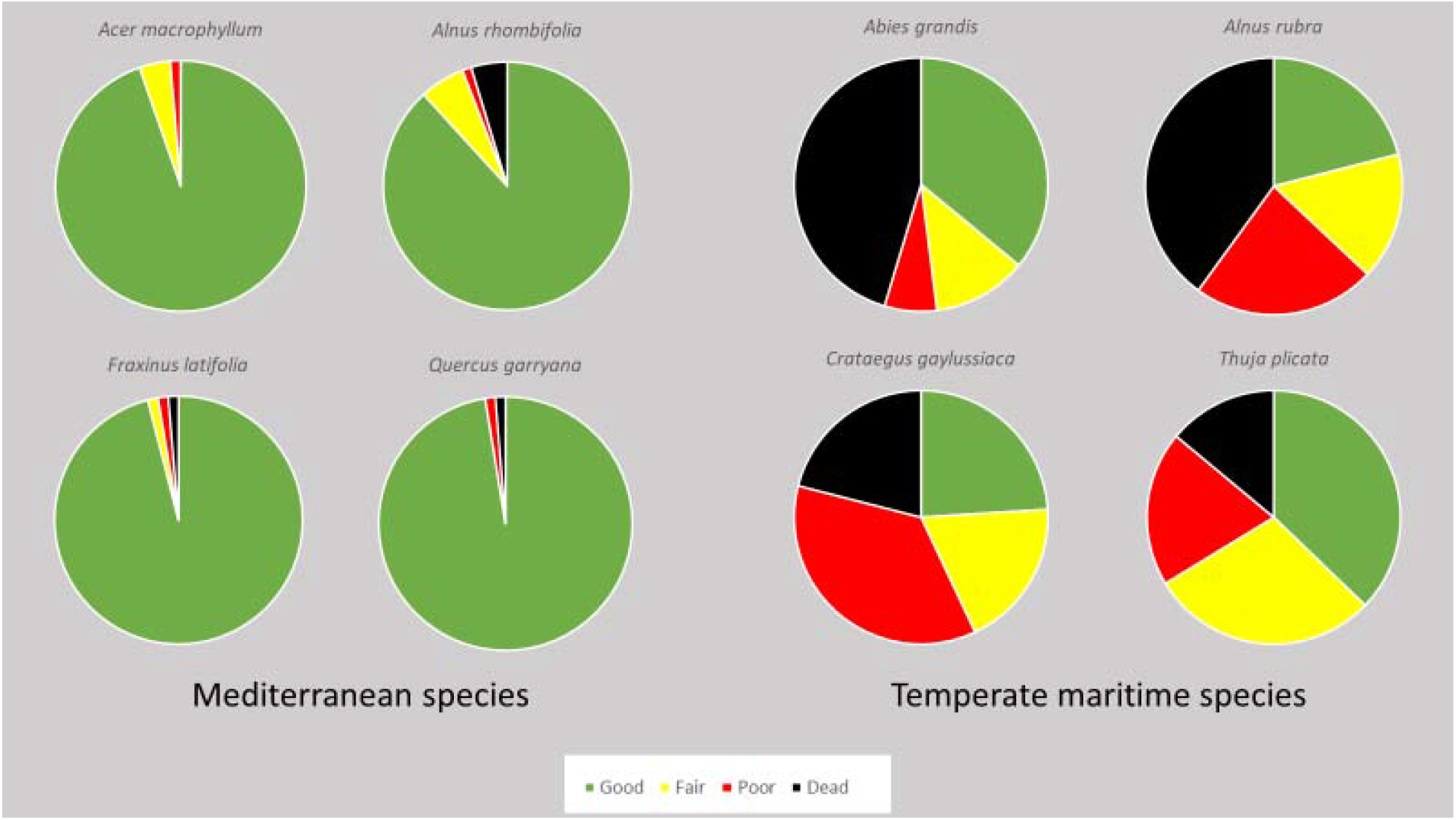
Comparison of Mediterranean (warm-adapted) and temperate maritime (cool-adapted) tree conditions in the Willamette Valley, Oregon

In each of the temperate maritime species, mortality was observed in all age and size classes, and no trends emerged in rates of mortality among the classes: mature trees, poles and saplings are all currently subject to high rates of mortality. All mortality noted in this study is estimated to have occurred within the past five years based on observations of mass tree mortality by foresters (Buhl 2016, Oregon Department of Forestry 2019, Oregon State University Extension Service, 2018) and the low degree of bark exfoliation and other characteristics of dead trees surveyed (class I snags only). This estimate is likely conservative based on Cline (1980). Cline estimates that the average time of transition from decay class I to decay class II occurs within 5 years post mortem in western Oregon Douglas-fir, which is a highly resinous tree with more durable bark relative to the four Pacific maritime species in this study. Additionally, Cline’s dataset only includes trees > 9cm DBH, while the current study includes trees as small as 5 cm diameter, which typically decay much faster than larger trees (Vanderwel 2006). We estimate that the observed 30% total mortality rate, therefore, equates to at least 6% annual mortality on average across cool-adapted species between 2015-2019 (table1).

### Pathogen Analysis in Black Hawthorn

Samples from 14 declining trees resulted in no one significant pathogen found in common among the plants assayed (Table 2). We did not recover any bacterial pathogens from the material (data not shown). The fungi recovered from tissues were largely saprophytic (*Cladosporium, Fusarium*), weak or opportunistic (*Epicoccum, Botrytis, Diaporthe*) or are pathogens often found in association with trees suffering from chronic stress (*Diplodia, Cytospora, Botryosphaeria*). The latter fungi can cause significant damage to affected plants, but are not considered capable of primary pathogenesis unless the plant is already compromised. The role of *Diaporthe eres* in hawthorn is unknown, although this fungus can be a pathogen to conifers and other woody plants.

**Table 2:** Sample dates, tissues examined, symptoms, and results of culturing from black hawthorn material showing decline symptoms

| Sample identifier | Date received | Tissue sampled | Symptoms | Outcome |
| --- | --- | --- | --- | --- |
| 17-1159 | 5 July 2017 | Branch | Dieback | *Diplodia* |
| 17-1492 | 15 August 2017 | Branch | Tip dieback | *Diaporthe* |
| 17-1516 | 16 August 2017 | Branch | Tip dieback, leaf spots | *Mycocentrospora acerina* (leaves); *Trichoderma* sp., *Phoma* sp. (branch) |
| 17-2058 | 2 November 2017 | Branch, root | Cankers, root necrosis, insect feeding injury | *Cytospora* sp. (branch, root) |
| 19-1693 | 2 October 2019 | Branch, root | Dieback | *Diaporthe eres* (branch) |
| 19-1775 | 10 October 2019 | Branch | Dieback | *Botryosphaeria* sp. |
| 19-1776 | 10 October 2019 | Branch | Dieback | *Cytospora* sp. |
| 19-1777 | 10 October 2019 | Branch | Dieback | *Botryosphaeria* sp. |
| 19-1778 | 10 October 2019 | Branch | Dieback | *Penicillium*, sterile fungi |
| 19-1802 | 17 October 2019 | Branch, bark | Tip dieback; bark borers (bark piece). | *Cytospora* sp., *Fusarium* sp (branch). |
| 19-1803 | 17 October 2019 | Branch, bark | Tip dieback; bark necrosis | *Diaporthe* sp. |
| 19-1804 | 17 October 2019 | Bark | Necrosis | Diverse sterile fungi |
| 19-1805 | 17 October 2019 | Branch, bark | Bark borers (bark pieces) | *Penicillium*, diverse sterile fungi (branch) |
| 19-1806 | 17 October 2019 | Branch, bark | Localized lesions, bark borers (bark). | *Epicoccum* sp., *Cladosporium* sp., *Botrytis* sp., *Fusarium* sp. |

## Discussion

The apparent absence of acute pathogen effects in black hawthorn mirrors findings of a previous investigation of declining alder in western Oregon, in which Sims (2015) found little evidence of acute pathogens, but rather several stress-associated insects and pathogens consistent with environmentally driven decline. Likewise, a recent advisory on dead and dying western redcedar by the Oregon Department of Forestry cites environmental stress as the cause of this decline rather than any acute pathogen (Oregon Department of Forestry 2019). The evidence presented here suggests environmental decline, and the data are clear that the decline is limited to cool-adapted trees. Decline of not just one, but four important tree species at their southerly and interior range margins, when coupled with climate data showing sharp increases in mean annual temperature and decreases in precipitation, provides compelling evidence of a broad, climate-related plant community shift, one with potential to affect the extent of some of the most productive forests in the world.

Abram et al (2016) trace the inception of detectable North American warming associated with land conversion and industrialization to the mid-19^th^ century. In the intervening ∼170-year period, there is no record of any change in the arborescent flora of the Willamette Valley. Now, after over a century and a half of increasing continental temperatures, several northerly arborescent species in the Willamette Valley are declining at rates that threaten their existence in this region in the near-term. If the rates of mortality estimated in this study continue, over half of the red alder and grand fir and a quarter of the black hawthorn and western redcedar present in the Willamette Valley in 2014 will be dead by 2025. If current temperature trends also continue their upward ascent, these species are likely very soon to cease to exist in this region. This outcome would corroborate projections from dynamic vegetation models showing substantial reductions in softwood in favor of hardwood/mixed forests (e.g., Hawkins et al., 2019; Sheehan et al., 2015; Turner et al., 2015) though those projections span a large of range outcomes due to uncertainties in future climate and model parameterizations (e.g., Hawkins et al., 2019; Shafer et al., 2015) Tree decline and mortality appear to be not just lagging indicators of climate change, but extremely delayed ones, suggesting the possibility of continued and accelerating emergence of costly and environmentally deleterious climate-driven range shifts in this region.

Most predicted plant community response to climate change is based on large-scale models of exceedingly complex systems. As Hamann (2006) concludes, “(if) currently observed climate trends continue or accelerate, major changes to management of natural resources will become necessary. Because of modeling uncertainties at small spatial scales, systematic field monitoring of biological response to climate change guided by our model predictions may be the best indicator for the need to implement management changes.” The simple approach presented here provides a means to rapidly assess local climate effects, and to inform species selection for nursery propagation and out-planting in the Willamette Valley. The same or similar methodologies can be replicated at any locality where significant climate change is occurring to assess its ecological effects and provide a basis for local land management decision making and on-the-ground action.

